# Heterologous expression of critical pathway genes leads to complex pattern of increased yield of bioplastic precursors in *Paraburkholderia sacchari* ITP 101

**DOI:** 10.1101/2024.11.21.624694

**Authors:** Dianna Morris, Niaz Bahar Chowdhury, Cheryl Immethun, Rajib Saha

## Abstract

Recent research endeavors have turned to sustainably generating useful chemicals from biological platforms. However, conventional model organisms, such as *Escherichia coli* and *Saccharomyces cerevisiae*, face limitations, particularly in terms of substrate range and yield for certain metabolites. In this study, we share our work toward the development of the non-model bacterium, *Paraburkholderia sacchari* (hereafter *P. sacchari*), as a microbial factory for the production of polyhydroxyalkanoates (PHAs), which are precursors for biodegradable plastic. The particular PHAs of interest produced by *P. sacchari* include poly(3-hydroxybutyrate) (PHB) and the co-polymer produced by the combination of PHB and 3-hydroxyvalerate (3HV) called poly(3-hydroxybutyrate-co-3-hydroxyvalerate) (PHBV). *P. sacchari* produces PHB from mixtures of hexose and pentose sugars commonly found in lignocellulosic biomass, however PHBV requires co-feeding with propionate. Both plastic precursors have industrial interest, so both PHB and 3HV were chosen as production targets. Due to studies in other bacteria demonstrating PHB yield can be improved by overexpressing genes for critical pathway enzymes, we hypothesized there is a bottleneck in the production pathway leading to PHB in *P. sacchari* as well. To explore this, heterologous genes coding for the three critical enzymes were taken from *Cupriavidus necator* H16 (hereafter *C. necator*) and inserted via plasmid; *phaA* and *bktb* (homologous genes for β-ketothiolase), *phaB* (acetyl-CoA reductase), and *phaC* (PHA polymerase). PHB production increased following overexpression of *phaB*, indicating acetoacetyl-CoA as the limiting enzyme. In fact, overexpression of *phaB* with the synthetic Anderson promoter, BBa_J23 104, increased titer by 162% over wildtype. On the other hand, strategies to improve 3HV had mixed results. Heterologous overexpression of propionyl-CoA transferase (*pct* from *C. necator*), which converts propionate into propionyl-CoA-the starting substrate for the 3HVproduction, showed a 145% increase in 3HV. Yet, internal sourcing of propionyl-CoA from succinyl-CoA following introduction of the sleeping beauty mutase (*sbm*) operon from *E. coli* showed no 3HV production. To this end, Max/Min Driving Force (MDF) thermodynamic analysis of critical PHBV pathways revealed two major limitations of 3HV production: 1) internal sourcing is not thermodynamically favorable; and 2) recycling of propionyl-CoA through the methyl citrate cycle (MCC) is more favorable than 3HV formation. Overall, we have shown promising progress and suggest future directions toward an industrially useful strain of *P. sacchari* for PHB and PHBV production.

**Graphical abstract:** 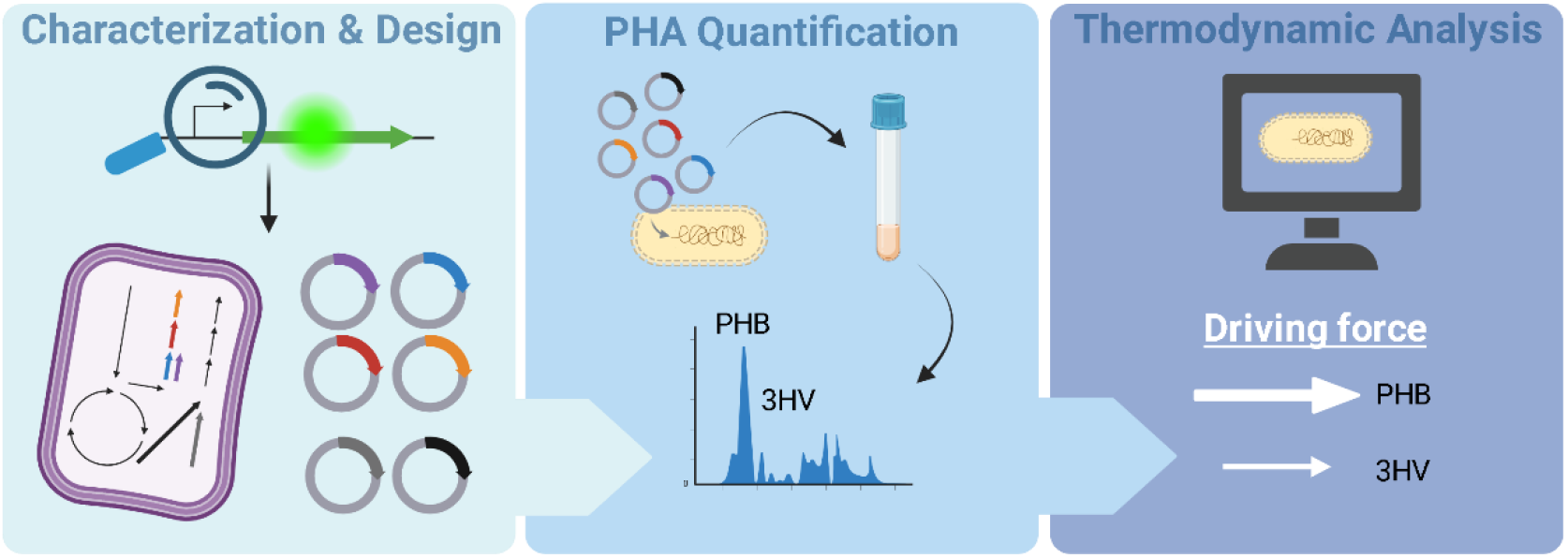

**Highlights:** *Paraburkholderia sacchari* has interest as a bioproduction platform for PHAs from complex feedstocks. Removal of PHA pathway bottleneck increases PHB yield by 162%. Improved conversion of fed-in propionate increases 3HV yield by 145%. Internal sourcing of propionyl-CoA does not successfully yield 3HV. Thermodynamic analysis provides insight into difficult conversion of propionyl-CoA to 3HV.

## 1. Introduction

The benefits of bioproduction (creation of a target chemical using a biological system) are numerous as it often provides a less toxic, highly specific, and sustainable option for generating a desired product (Hudiburg et al., 2016). Plant biomass can serve as a renewable and reliable source of carbon for energy generation and product synthesis (Robertson et al., 2017). Through bioproduction, products such as the bioplastics, polyhydroxyalkanoates (hereafter **PHAs**), a biodegradable alternative to petroleum-based plastics, could be produced sustainably, both economically and environmentally. Non-model organisms, though difficult to work with from lack of characterization, expand the range of bioproduction due to native capabilities (Fatma et al., 2020). One such non-model microbe is *Paraburkholderia sacchari* IPT101 (hereafter, *P. sacchari*); a Gram-negative bacterium isolated from sugarcane soil that can accumulate up to 75% of its cell dry weight as PHAs (Gomez et al., 1996). It has benefits over other potential PHA producers, such as *Cupriavidus necator* H16, due to a fast specific growth rate (Gomez et al., 1996) and, most importantly, a broad range of usable carbon sources that can be converted into PHAs including glucose, xylose, fructose, arabinose, mannose, sucrose, and galactose (Brämer et al., 2001; Dietrich et al., 2018). The particular PHAs of interest produced by *P. sacchari* include poly(3-hydroxybutyrate) (hereafter PHB) and the co-polymer produced by the combination of PHB and 3-hydroxyvalerate (hereafter 3HV) called (poly(3-hydroxybutyrate-co-3-hydroxyvalerate) (hereafter PHBV) (Gomez et al., 1996). PHB is the main monomer produced, yet PHBV is particularly interesting due to its improved toughness and elasticity (Liu et al., 2014). Synthesis of both PHB and 3HV follow a three-step pathway using the enzymes β-ketothiolase, acetoacetyl-CoA, and PHA synthase (Alvarez-Santullano et al., 2021) (shown in Figure 1). The starting substrate for PHB is acetyl-CoA, whereas the starting substrate for 3HV is propionyl-CoA.

**Figure 1.**
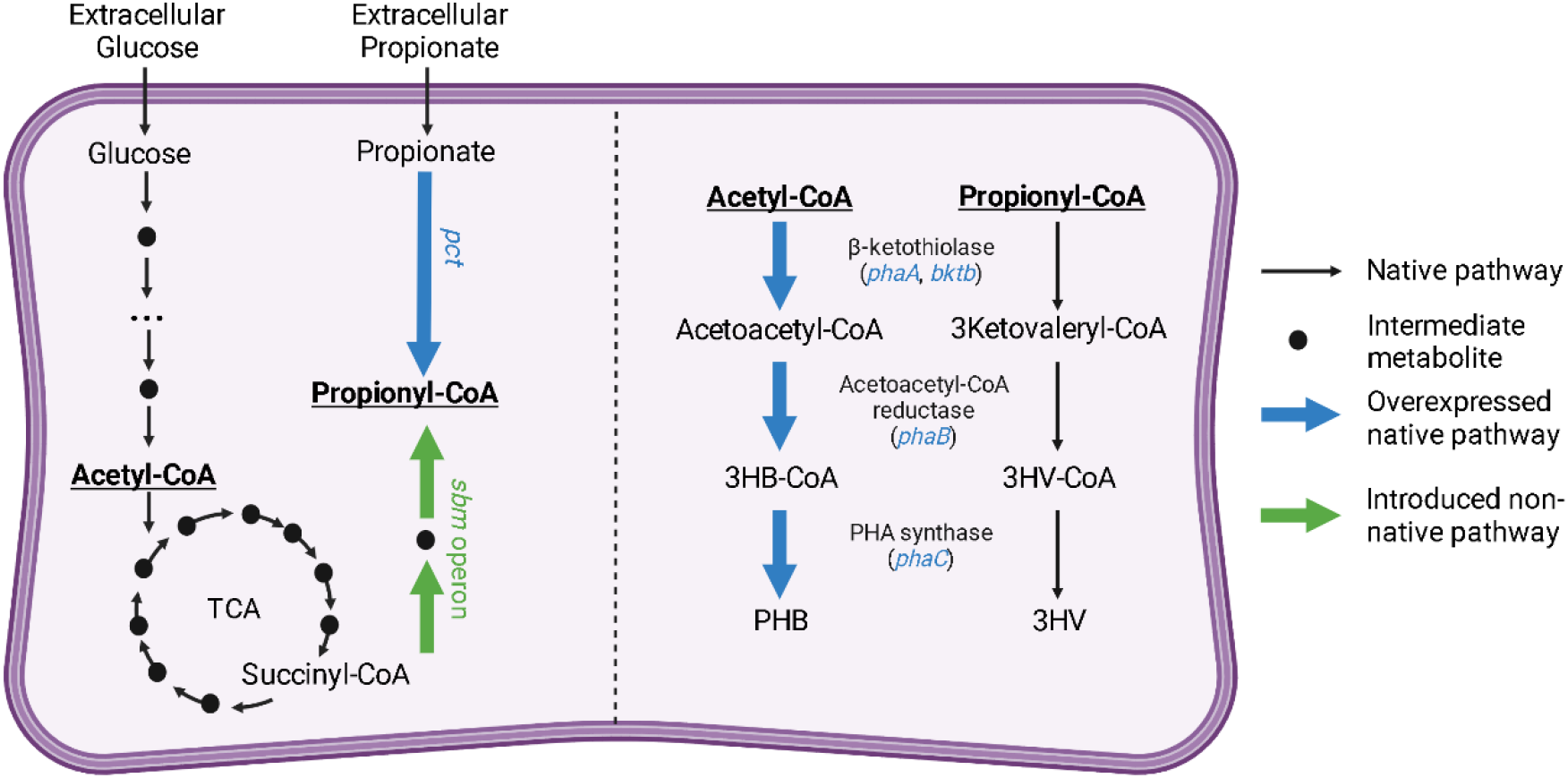
Simplified metabolic pathways of *P. sacchari* associated with PHB and 3HV production and planned genetic targets.

Previous work has laid groundwork for metabolic engineering of this bacterium to improve the yields of PHAs, including genome sequencing (Alexandrino et al., 2015), UV irradiation mutant characterization (Brämer et al., 2002; Lopes et al., 2011; Silva et al., 2000), introduction of heterologous operons (Mendonça et al., 2017), overexpression of transcriptional regulators (Guamán et al., 2018), and gene knockout via homologous recombination (Pereira et al., 2009). PHB has been increased through metabolic engineering that targeted carbon catabolite repression (CCR) through characterization of xylose metabolism(Lopes et al., 2009) and overexpression of critical pathway genes (Guamán et al., 2018; Guaman et al., 2018). 3HV has also been increased by creation of a UV mutant (Silva et al., 2000) which has been shown to have a modified 2-methylcitric acid cycle (MCC) (Brämer et al., 2002). To recreate the genetic modifications found in the mutant, two key genes in the MCC pathway (*prpC* encoding 2-methylcitrate synthase and *acnM* encoding aconitate hydratase) were knocked out via homologous recombination and demonstrated increased yield over wildtype, however did not reach levels of the UV mutant strain (Pereira et al., 2009). While these works have shown promise in developing *P. sacchari* as a bioproduction platform for PHB and PHBV synthesis, here we share application of additional approaches, an overview of which can be seen in Figure 1.

In engineered *E. coli*, the metabolic limitations of the PHB synthesis pathway has been explored and overexpression of PHB pathway genes identified acetoacetyl-coA reductase (*phaB*) as a bottleneck (Tyo et al., 2010). While this has not been replicated in *P. sacchari*, we hypothesize that such a bottleneck is also present in this non-model, and that overexpression of the *phaB* gene improves PHB production. Growth studies have been performed that show PHBV production can only be achieved with additional co-fed substrates such as propionate. When propionate is added to the media, *P. sacchari* produces PHBV with a 3HV fraction of 28%, however, this is also accompanied by a drop in overall PHA accumulation (45%) and inhibition to growth (Mendonça et al., 2014). This demonstrates a limitation in how much propionate can be converted to 3HV. To address this limitation, we can look at two strategies previously used in other bacterial species: 1) upregulation of propionyl-CoA transferase (gene *pct*), the enzyme which converts proprionate and acetyl-CoA into propionyl-CoA, a strategy previously used in *E. coli* (Jung et al., 2019); and 2) internal sourcing of propionyl-coA from succinyl-coA by introduction of the sleeping beauty mutase (*sbm*) operon containing genes *sbm* and *ygfG* encoding a novel (2R)-methylmalonyl-CoA mutase and a (2R)-methylmalonyl-CoA decarboxylase respectively, a strategy previously used in *Salmonella enterica* (Aldor et al., 2002). We hypothesize that upregulation of *pct* increases 3HV by allowing for more efficient uptake of fed-in propionate and that introduction of the *sbm* operon allows for 3HV production even in the absence of fed-in propionate. Due to the non-model nature of this bacterium, we cannot assume that strategies to increase PHB or 3HV yield in other bacteria will be directly transferable. Therefore, we also report on the development of a thermodynamic model to examine overall pathway driving force and associated thermodynamic bottlenecks of the pathways investigated in this paper.

## 2. Materials/Methods

### 2.1 Strain/growth information

*Paraburkholderia sacchari* IPT101 (Brämer et al., 2001) (previously classified as *Burkholderia* (Dobritsa and Samadpour, 2016)) seed cultures were started from frozen stock inoculated into 4 mL mineral media (MM)(Rocha et al., 2008) with 3 gL^-1^ Na_2_HPO_4_ and 30 gL^-1^ glucose as the carbon source in 14 mL Falcon™ round-bottom polystyrene tubes and incubated for 48 hours at 30° C and 250 rpm constant shaking. *Cupriavidus necator* H16 and *E. coli* DH10B were both started from frozen stock and cultured in luria broth (LB) media at 30° C and 250 rpm constant shaking for 24 hours before genomic DNA (gDNA) harvesting. *E. coli* DH10B was used for plasmid construction and maintenance. Kanamycin (50µgL^-1^) was added for plasmid maintenance in both *E. coli* and *P. sacchari* cultures when appropriate.

### 2.2 Part construction and transformation

To construct the plasmids used in this study, primers were purchased from Eurofins Genomics (Louisville, KY), PCR templates were amplified with Phusion Hot Start II High-Fidelity DNA Polymerase (Thermo Scientific) and amplicon purification was done with either PureLink Gel Extraction Kit (Invitrogen, Carlsbad, CA) or PureLink Quick PCR Purification Kit (Invitrogen). Genomic DNA (gDNA) used in this study was obtained with Zymo Research Quick-DNA Fungal/Bacterial Miniprep Kit. Strains and plasmids created in this study, sequences of genetic parts, and primers used to amplify target sequences are listed in Table 1, and Supplementary Tables 1 and 2 respectively.

**Table 1:** Description and source of strains and plasmids used in this study.

| Strain | Description | Source or Reference |
| --- | --- | --- |
| ***Paraburkholderia sacchari* IPT 101** | Wildtype | (Brämer et al., 2001) |
| ***Cupriavidus necator* H16** | Wildtype | DSM 428 |
| ***E. coli* DH10B** | T1 phage resistant and endonuclease I (endA1) deficient | New England Biolabs® |
| **sPS001** | *P. sacchari* containing SSBIO-PS001 | This study |
| **sPS002** | *P. sacchari* containing SSBIO-PS002 | This study |
| **sPS003** | *P. sacchari* containing SSBIO-PS003 | This study |
| **sPS004** | *P. sacchari* containing SSBIO-PS004 | This study |
| **sPS005** | *P. sacchari* containing SSBIO-PS005 | This study |
| **sPS006** | *P. sacchari* containing SSBIO-PS006 | This study |
| **sPS007** | *P. sacchari* containing SSBIO-PS007 | This study |
| **sPS008** | *P. sacchari* containing SSBIO-PS008 | This study |
| **sPS009** | *P. sacchari* containing SSBIO-PS009 | This study |
| **sPS010** | *P. sacchari* containing SSBIO-PS010 | This study |
| **sPS011** | *P. sacchari* containing SSBIO-PS011 | This study |
| **sPS012** | *P. sacchari* containing SSBIO-PS012 | This study |
| **sPS013** | *P. sacchari* containing SSBIO-PS013 | This study |
| **sPS014** | *P. sacchari* containing SSBIO-PS014 | This study |
| **sPS015** | *P. sacchari* containing SSBIO-PS015 | This study |
| **Plasmid** |  |  |
| **pBBR1MCS-2** | Expression vector containing the pBBR1 origin of replication, *lacZ* under the control of pLAC, and kanamycin resistance | Addgene 85168 (Watertown, MA)(Kovach et al., 1995) |
| **SSBIO-PS001** | *gfpUV* under the control of pLAC on pBBR1MCS-2 backbone with kanamycin resistance | This study |
| **SSBIO-PS002** | *gfpUV* under the control of BBa_J23 100 on pBBR1MCS-2 backbone with kanamycin resistance | This study |
| **SSBIO-PS003** | *gfpUV* under the control of Bba_J23 104 on PBBR1MCS-2 backbone with kanamycin resistance | This study |
| **SSBIO-PS004** | *gfpUV* under the control of Bba_J23 107 on PBBR1MCS-2 backbone with kanamycin resistance | This study |
| **SSBIO-PS005** | *gfpUV* under the control of Bba_J23 115 on PBBR1MCS-2 backbone with kanamycin resistance | This study |
| **SSBIO-PS006** | *phaA* under the control of pLAC on PBBR1MCS-2 backbone with kanamycin resistance | This study |
| **SSBIO-PS007** | *bktb* under the control of pLAC on PBBR1MCS-2 backbone with kanamycin resistance | This study |
| **SSBIO-PS008** | *phaB* under the control of pLAC on PBBR1MCS-2 backbone with kanamycin resistance | This study |
| **SSBIO-PS009** | *phaC* under the control of pLAC on PBBR1MCS-2 backbone with kanamycin resistance | This study |
| **SSBIO-PS010** | *phaA* under the control of Bba_J23 104 on PBBR1MCS-2 backbone with kanamycin resistance | This study |
| **SSBIO-PS011** | *bktb* under the control of Bba_J23 104 on PBBR1MCS-2 backbone with kanamycin resistance | This study |
| **SSBIO-PS012** | *phaB* under the control of Bba_J23 104 on PBBR1MCS-2 backbone with kanamycin resistance | This study |
| **SSBIO-PS013** | *phaC* under the control of Bba_J23 104 on PBBR1MCS-2 backbone with kanamycin resistance | This study |
| **SSBIO-PS014** | *pct* under the control of pLAC on PBBR1MCS-2 backbone with kanamycin resistance | This study |
| **SSBIO-PS015** | *sbm* operon under the control of pLAC on PBBR1MCS-2 backbone with kanamycin resistance | This study |
| **AS-Plac(c)-GFPuv-amp** | *gfpUV* under control of pLAC, rrnT1 terminator, EC ori, AS ori, with kanamycin resistance | Unpublished, Saha Lab (Lincoln, NE) |

Plasmid SSBIO-PS001 was created using Hot Fusion (Fu et al., 2014). The plasmid backbone pBBR1MCS-2 (Kovach et al., 1995)(purchased from Addgene 85168), excluding the *LacZ* gene, was linearized to obtain the PBBR1 origin of replication, kanamycin resistance, and the promoter pLAC. In place of *LacZ*, the *gFPuv* gene (originally from plasmid pBbB7a-GFP (Lee et al., 2011) purchased from Addgene 35358, Watertown, MA) was amplified along with the rrnBT1 terminator(Mutalik et al., 2013) from a plasmid constructed from a previous project AS-Plac(c)-GFPuv-amp (unpublished, Saha Lab at UNL). Also using the hot fusion method, plasmid SSBIO-PS001 was linearized excluding the gFPuv gene and *phaA, bktb, phaB, phaC, pct*, and the *sbm* operon were inserted after amplification from gDNA (all from *C. necator* except *sbm* from *E. coli*) to create plasmids SSBIO-PS006-9 and SSBIOPS014 & 15 respectively. Plasmids SSBIO-PS002-5 were created using blunt end ligation to replace the pLAC promoter with four Anderson promoter variants; BBa_J23 100, BBa_J23 104, BBa_J23 107, and BBa_J23 115 (Anderson, 2008) respectively. The plasmid SSBIO-PS001 was linearized, excluding the pLAC promoter, using primer pairs that contained tails with Anderson promoter sequences. The resulting PCR products were treated with DpnI (Thermo Fisher Scientific, Waltham, MA), gel-extracted, phosphorylated using T4 PNK (Thermo Fisher Scientific), and assembled into plasmids using T4 DNA ligase (Thermo Fisher Scientific), following the guidelines for the self-circularization of linear DNA. Following the same procedure, plasmids SSBIO-PS010-13 were created using primer pairs containing the Anderson promoter BBa_J23 104 to linearize SSBIO-PS006-9 respectively.

Plasmids were obtained from cultures of *E. coli* DH10B using the PureLink Quick Plasmid Miniprep Kit (Invitrogen) and were sequence-verified. These plasmids were introduced into *P. sacchari* via a room-temperature electroporation method (Tu et al., 2016). Strains sPS001-sPS015 were then stored at −80°C in 40% glycerol for further analysis.

### 2.3 Characterization of constitutive promoters

Seed cultures of sPS001-sPS005 along with wildtype *P. sacchari* (hereafter WT) were diluted to 0.1 OD_600_ in fresh MM and grown an additional 24 hours in biological triplicate. 500 µL of each culture was then collected, pelleted, and washed twice in phosphate buffer saline (PBS) before being resuspended in a final volume of 200 µL PBS. Fluorescence (excitation 395 nm and emission 509 nm) was collected using a Molecular Devices SpectraMax i3x Multi-Mode Microplate Detection Platform. Results were processed by subtracting background WT and PBS fluorescence and normalized to cell density (absorbance 600 nm).

### 2.4 Nitrogen starvation, extraction of PHAs and HPLC measurements

To assess PHA yield in the engineered strains, a nitrogen starvation method was used. Seed cultures of sPS006-15 along with WT were diluted, in biological triplicate, to 0.1 OD_600_ in fresh MM, grown an additional 20 hours to reach late exponential phase, pelleted, and resuspended in fresh MM excluding Na_2_HPO_4_ for nitrogen starvation. WT and strain sPS014 were grown both in the absence and presence (1 gL^-1^) of sodium propionate (Sigma-Aldrich). Samples were grown an additional 24 hours, pelleted, and stored at −80°C for further processing. PHA extraction was done using an alkaline digestion high performance liquid chromatography (HPLC) method (Satoh et al., 2016).

The cell pellets resuspending in 1 mL water were used as the starting material. Measurements were taken following the published method on an Agilent 1100 HPLC (Agilent Technologies, Santa Clara, CA, USA) using an Eclipse XDB-C18 column (Agilent) and final results were divided by 4 to account for the starting culture volume of 4 mL.

### 2.5 Transmission electron microscopy (TEM)

To visualize PHA produced by WT and sPS006-15 strains grown under the described nitrogen starved conditions (section 2.4) were prepared for transmission electron microscopy (TEM) according to an established method (Graham and Orenstein, 2007). Cells were fixed in a solution of 2% glutaraldehyde and 1.5% paraformaldehyde in 100 mM sodium cacodylate buffer for over 1 hour at room temperature, followed by incubation at 4°C overnight. The samples were then washed three times with sodium cacodylate buffer and post-fixed in 1% osmium tetroxide in deionized water for 1 hour at room temperature. After two washes in water, the samples were dehydrated through a graded ethanol series and embedded in Spurr medium using standard TEM processing techniques. Ultrathin sections were prepared using a Leica UC7 ultramicrotome, then stained with 1% uranyl acetate and 1% lead citrate. Images were acquired with a Hitachi H7800 with a side-mount camera (16BM) at the Microscopy Core Research Facility, Center for Biotechnology, University of Nebraska-Lincoln and processed using ImageJ (Schneider et al., 2012).

### 2.6 Model construction and Max/Min Driving Force

We next used the Max/Min Driving Force (MDF) analysis to assess thermodynamic drive and bottlenecks (Noor et al., 2014). Since the *P. sacchari* genome was not yet in the KEGG Database (Kanehisa and Goto, 2000), the available genome of a closely related and PHA producing organism, *Paraburkholderia xenovorans* (Alvarez-Santullano et al., 2021; Goris et al., 2004; Sawana et al., 2014), was used to generate a list of genes involved in glycolysis, the TCA cycle, and propanoate metabolism. Known genes in *P. sacchari* for PHB and 3HV metabolism (Alexandrino et al., 2015; Alvarez-Santullano et al., 2021) were added in as well. Reactions associated with the genes were found using the Metabolic Atlas (Li et al., 2023a) and Equilibrator (Beber et al., 2022) was used to estimate the change in Gibbs Free Energy (Δ_r_G^’0^) for each reaction in the proper direction. The final polymerizing reaction for both PHB and 3HV were not included due to difficulty of estimated Δ_r_G^’0^ for such reactions (Bley and Dzubiella, 2022; Panayiotou, 2024), and the thermodynamic feasibility in the forward direction is assumed. The constructed model was used to run MDF according to the following formulation:

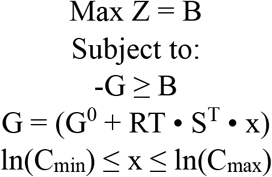

Where B is the driving force, G is the calculated Gibbs free energy, G^0^ is the standard change in Gibbs free energy estimated from Equilibrator (Beber et al., 2022), R is the gas constant, T is the temperature-in this instance 30° C (303.15° K) to mimic growth conditions, S^T^ is the stoichiometric matrix of all the reactions and metabolites, x is the metabolite concentration, and C_min_ and C_max_ denote the allowed range of metabolite concentration. Constraints for ATP/ADP concentration ratio was set to atleast 10 and for NADH/NAD to at least 0.1 (Noor et al., 2014). Since metabolite concentrations have not been measured in *P. sacchari*, for the other metabolite, the concentration range was set from 1 µM to 100 mM based on ranges determined in *E. coli* (Bennett et al., 2009). The objective value, Z, returns the lowest calculated driving force, representing the reaction with the least negative calculated ΔG. A positive Z value indicates thermodynamic feasibility of the entire pathway.

## 3. Results and Discussion

### 3.1 Characterization of constitutive promoters

Since pLAC is a commonly used promoter with the potential to be included in an inducible system and has been previously used in *P. sacchari* (Guamán et al., 2018), this was included in our promoter set. The Anderson promoters, synthetic promoters developed for use in *E. coli*, were also included. These were designed to provide a range of possible expression based on how closely they match the sigma factor consensus sequence of *E. coli*’s main RNA polymerase (Anderson, 2008) yet have also been shown to allow for a range of expression several non-model bacteria including *Pseudomonas putida* (Pearson Allison et al., 2023), *Chromobacterium violaceum* (Liow et al., 2019), and *Actinobacillus succinogenes* (Long et al., 2021). A subset of four promoters spread across the range of expression in *E. coli* were chosen for testing: BBa_J23 100 (strength: 1), BBa_J23 104 (strength: 0.72), BBa_J23 107 (strength: 036), and BBaJ23_115 (strength: 0.15). The construction of the plasmids and strain IDs is shown in Figure 2A. Fluorescence measurements indicate that all five tested promoters in sPS001-5 demonstrate expression in *P. sacchari* as can be seen in Figure 2B. Student’s t-tests indicated that all the Anderson promoters showed significantly greater expression than the pLAC promoter in sPS001 (p<0.05) and the BBa_J23 104 promoter in sPS003 showed significantly greater expression than all other promoters (p<0.01). The measured expression of the Anderson promoters does not match the pattern of decreasing strength seen in *E. coli* (Anderson, 2008). This is not surprising, as sigma factors responsible for RNA polymerase binding are known to vary between bacterial species (Browning and Busby, 2004) and the pattern of expression across the range of promoters has been seen to be different in each bacteria they have been tested in (Liow et al., 2019; Long et al., 2021; Pearson Allison et al., 2023). This mismatching pattern suggests that *P. sacchari* has a slightly different consensus sequence than *E. coli*.. However, the Anderson promoters can still be used in *P. sacchari* to provide options for tuned constitutive gene expression. For purposes of this study, pLAC and BBa_J23 104 were used for plasmid design to compare two levels of expression of metabolic genes.

**Figure 2.**
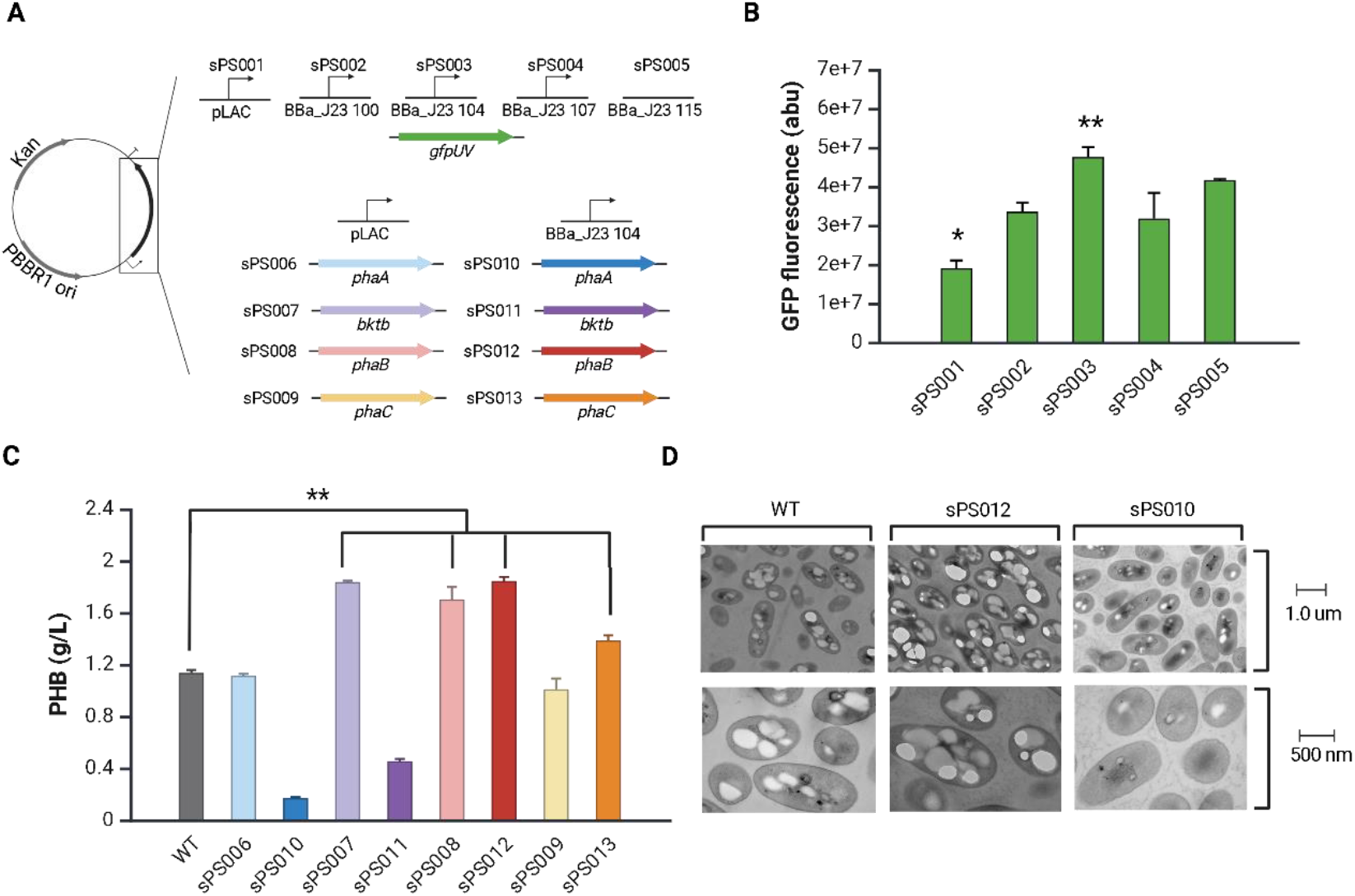
Design and results for characterization of promoters and overexpression of PHA pathway genes. A) Diagram illustrating genetic parts transformed into each strain referenced in this figure. B) Graph representing normalized fluorescence (abu) for strains sPS001-sPS005. C) Graph representing quantified PHB (g/L) in Wildtype *P. sacchari* (WT) and strains sPS006-sPS013. D) TEM images for WT, sPS012, and sPS010 at 20x magnification and 50x magnification. White inclusions are PHB. All error bars represent standard deviation, and relevant significant differences are indicated as follows: *: p < 0.05, **: p < 0.01.

### 3.2 Overexpression of PHB pathway genes

To overexpress PHB pathway genes, three genes from C. necator (*phaA, phaB*, and *phaC*) were chosen due to their proven ability to convey PHB production capabilities to natively non-producing bacteria (Horng et al., 2011). An additional homolog of *phaA, bktb*, was also included as it has been shown to be important for β-ketothiolase activity in *C. necator* (Lindenkamp et al., 2010). The design of the plasmids constructed, and the strain numbers can be seen in Figure 2A. When each of these genes was inserted into *P. sacchari* under the control of pLAC (strains sPS006-sPS009), both *phaB* (sPS008) and *bktb* (sPS007) conveyed statistically significant increases in PHB yield over WT (student’s t-test: p<0.01 and p<0.001 respectively) as can be seen in Figure 2C. Overexpression of *phaA* and *phaC*, (strains sPS006 and sPS009 respectively), showed no significant difference from WT (p>0.05). However, when the same genes were inserted under the control of the stronger promoter, BBa_J23 104, the pattern changes with *phaA* (sPS010) and *bktb* (sPS011) showing significantly lower PHB yield and *phaC* (sPS013) showing significantly higher yield (p<0.001). This increased overexpression of *phaB* (sPS012) continued to show increased yield over WT (p<0.001) although not at a level higher than strain sPS008 (p = 0.07). TEM results support these findings by showing the visual difference in amount of PHB inclusions (white masses) across the different conditions. Images for WT, the significantly increased amount in sPS012, and the significantly decreased amount in sPS010 are shown in Figure 2D.

Several inferences could be gathered from these data. First, overexpression of *phaB* consistently increases PHB production, suggesting that the enzyme acetoacetyl-coA reductase acts as a bottleneck along the PHB pathway in *P. sacchari*. This finding is supported by previous studies that observe this same pattern (Tyo et al., 2010) and kinetic studies that showed low catalytic efficiency of this enzyme (Sim et al., 1997; Wang and Lee, 1997). Second, the data suggest that the β-ketothiolases encoded by *phaA* and *bktb* play unique roles in PHB production. It has been shown that there are 14 homologs for β-ketothiolase in *C. necator* (Pohlmann et al., 2006) and mutant studies have identified differing outcomes depending on which homolog is knocked out (Lindenkamp et al., 2010). It has been noted that *bktb* is required for 3HV production in *C. necator*, however, it has not been previously demonstrated to play a more important role in PHB production than *phaA*. This result points to a need to identify the homologs that may exist in *P. sacchari* to determine which homolog plays a more crucial role in PHB production. Third, the decrease in production following the increased overexpression of *phaA* and *bktb* suggests too much β-ketothiolase limiting the PHB pathway. This is supported by previous work showing that β-ketothiolase is thermodynamically unfavorable in the forward direction and that too much enzyme could increase flux in the backward direction (Alsiyabi et al., 2021; Tyo et al., 2010). Lastly, the increase of PHB following overexpression of *phaC*, but only in the higher expression condition, suggests that PHA synthase is not a bottleneck of the pathway, but in high concentrations can pull flux toward PHB production. This could be explained by overexpression of *phaC* having a “pull” effect on the PHB pathway where overexpression of a terminal enzyme can increase overall flux by increasing the rate of consumption of the final substrate(Volk et al., 2023). This effect can be seen in other cases, such as in synthesis of terpenes in *Synechocystis* PCC 6803, where overexpression of the terminal enzyme, isoprene synthase, increased isoprene production (Englund et al., 2018). Overall, these results show the complexities of proper enzyme balance in the PHB pathway, yet provide a clear demonstration that removing the β-ketothiolase bottleneck is a successful strategy for increasing PHB production in *P. sacchari*.

### 3.3 Propionyl-coA sourcing strategies

The *pct* gene from *C. necator* was chosen for overexpression since it was shown to generate heterologous overexpression of propionyl-CoA transferase in previous studies (Jung et al., 2019) and the *sbm* operon from *E. coli* was chosen as it has been shown to convey the ability to convert succinyl-CoA to propionyl-CoA in non-native species (Aldor et al., 2002) Both act as strategies to increase intracellular propionyl-CoA, which is the starting metabolite in the 3HV pathway. The schematic showing the insertion of genes *pct* and the *sbm* operon onto the plasmid vector and the corresponding strain IDs can be seen in Figure 3A.

**Figure 3.**
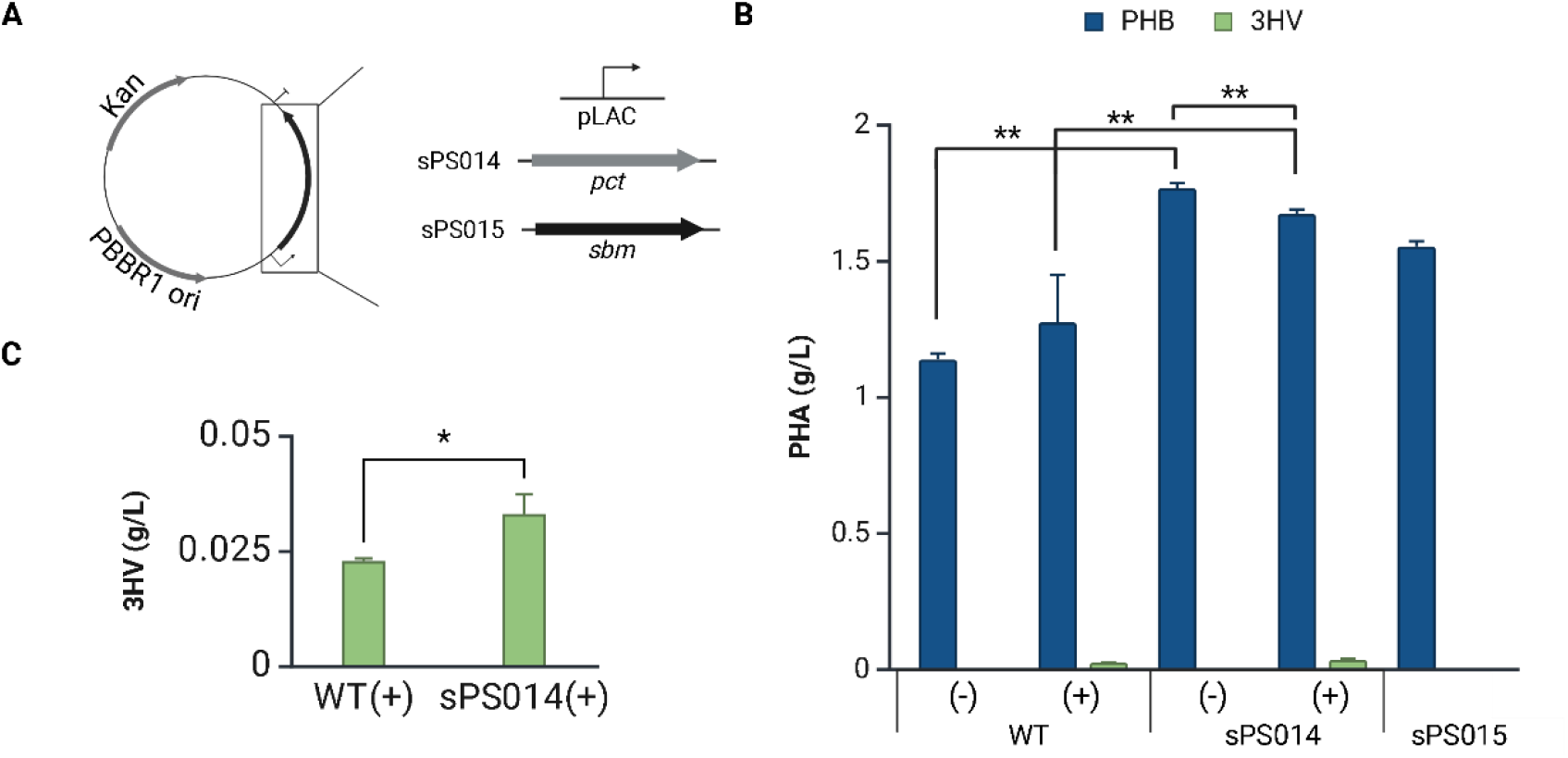
Design and results for overexpression and introduction of genes to target propionyl-CoA sourcing. A) Diagram illustrating genetic parts transformed into each strain referenced in this figure. B) Graph representing quantified PHA (g/L) in Wildtype *P. sacchari* (WT) and strains sPS014 and sPS015. For WT and sPS014, (-) and (+) indicated the absence or presence (1 g/L) of propionate respectively. C) A blown up graph showing 3HV yield in WT and sPS014 in the presence of propionate. All error bars represent standard deviation, and relevant significant differences are indicated as follows: *: p < 0.05, **: p < 0.01.

Quantified PHB and 3HV results are shown in Figure 3B and a closer look at the 3HV results can be seen in Figure 3C. The addition of propionate does result in 3HV production in the WT strain, as expected. Results also show no significant increase in PHB yield following addition of propionate (student’s t-test: p = 0.13). In strain sPS014, which has overexpressed *pct*, 3HV is again observed after the addition of propionate.

Notably, there is a significant difference in PHB yield; the addition of propionate leads to a slight decrease in yield (p < 0.01). As hypothesized, when comparing WT and sPS014, results show an increase in 3HV yield in the presence of propionate (p < 0.01) and, interestingly, also show an increase in PHB yield in both the absence and presence of propionate (p < 0.01). These data suggest that overexpressing the *pct* gene enhances the intracellular synthesis of propionyl-CoA, thereby increasing the available starting material for 3HV production when propionate is present. However, the higher yield of PHB in strain sPS014, even without propionate, indicates that the overexpression of *pct*—and the presumed upregulation of propionyl-CoA transferase—may also be activating another reaction that contributes to the increased PHB production. In fact, the decrease in PHB in the presence of propionate in sPS014 suggests the consumption of propionyl-CoA transferase acting on propionate is in competition with the unidentified reaction. Previous studies have shown that propionyl-CoA transferase can act on substrates other than propionate including malonyl-CoA (Li et al., 2023b) and lactic acid (Shi et al., 2022). Our findings further suggest the promiscuous activity of this enzyme. Additional exploration would be needed to identify the precise metabolic activity resulting in increased PHB yield.

Following the introduction of the *sbm* operon (strain sPS015), results show no 3HV yield, yet do show a significant increase in PHB yield compared to WT (p < 0.001). The 3HV results alone make it appear that internal sourcing of succinyl-CoA to propionyl-CoA is not successful. However, the increase of PHB suggests that introduction of *sbm* is allowing for internal sourcing of succinyl-CoA that is then being directed toward PHB synthesis rather than 3HV synthesis. This is possible if the resulting propionyl-CoA is going through the methyl citrate cycle (MCC) and being recycled back to pyruvate and subsequently acetyl-CoA, which serves as the PHB precursor. This finding is supported by previous work suggesting the MCC is efficient at recycling propionyl-CoA back to pyruvate and showing that elimination of critical genes in the MCC pathway increases 3HV yield (Brämer et al., 2002; Pereira et al., 2009). While it is not unexpected that there would be some increase in PHB following introduction of the *sbm* operon, it is somewhat surprising to see no 3HV production whatsoever.

### 3.4 MDF analysis to explore 3HV and PHB results

To further explore the results showing lack of 3HV production in sPS015 and increased PHB production in sPS014 and sPS015, MDF analysis was conducted in this study. MDF analysis optimizes the total driving force of a given pathway while considering the biologically relevant concentrations of various metabolites (Noor et al., 2014). MDF analysis also accepts estimated ΔGs and does not require measured enzyme activity; data that has not been collected in *P. sacchari*. Therefore, this analysis was chosen to look for thermodynamic bottlenecks and compare driving forces that may impact the fate of key metabolites in PHB and 3HV production.

The *P. sacchari* MDF model consisted of 35 reactions across glycolysis, the tricarboxylic acid (TCA) cycle, propionate metabolism including the methyl citrate cycle (MCC), and polyhydroxyalkanoate (PHA) metabolism (shown in Figure 4A) and it returns an overall positive driving force of the entire pathway, 0.01364 KJ/mol, indicating the pathway is thermodynamically feasible with the lowest calculated ΔG being −0.01364 KJ/mol. Reactions with this lowest value are indicated in red on Figure 4A and the non-native pathway introduced with the *sbm* operon in sPS015 is shown in green. The cumulative ΔGs across the pathway were compared using the estimates for each reaction (before MDF) and the calculated values using MDF (after MDF) as shown in Figure 4B. The values calculated for each reaction using MDF were more negative and the overall cumulative value much lower than the value based on the estimates. This demonstrates that when substrate concentration is considered across the entire pathway as a whole with MDF, the thermodynamic drive can be seen to be greater than what would be estimated when looking at each reaction individually. This more realistic and wholistic view of the pathway gives more contextually accurate calculated ΔGs.

**Figure 4.**
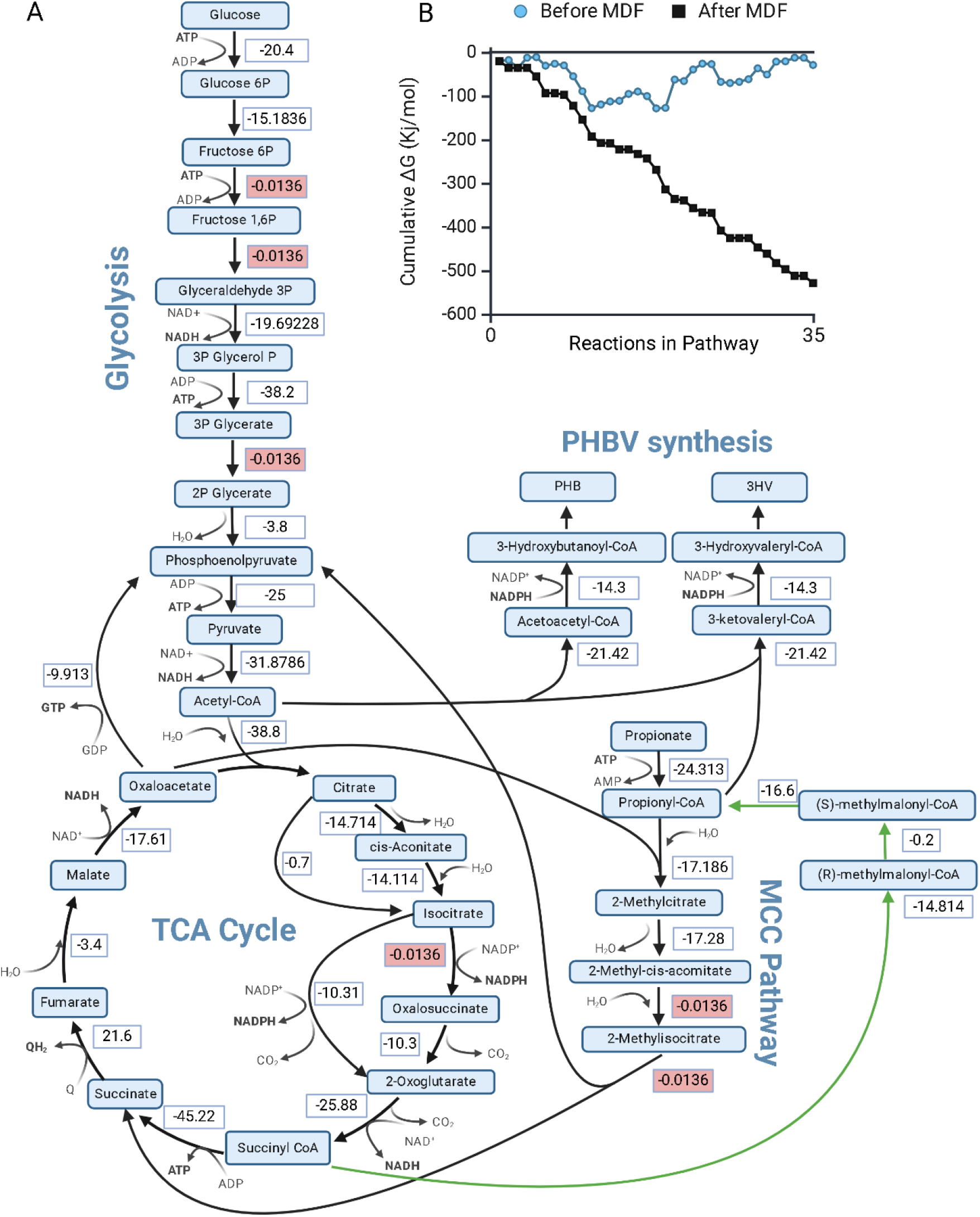
MDF model results. A) Core *P. sacchari* model with calculated ΔGs included. Values in red boxes represent bottleneck reactions. Green arrows indicate the non-native pathway introduced in sPS015 B) Cumulative ΔG values both before and after MDF analysis.

The calculated ΔGs were used to assess the fate of succinyl-CoA and propionyl-CoA. For succinyl-CoA, the cumulative ΔG going to propionyl-CoA formation through the novel *sbm* operon pathway is −31.614 KJ/mol, whereas going to acetyl-CoA formation through the TCA cycle is −154.6216 KJ/mol. This comparison suggests that despite the introduction of genes that could pull succinyl-CoA to propionyl-CoA and create potential for increased 3HV production, there is more thermodynamic drive keeping succinyl-CoA in central metabolism and recycling back to acetyl-CoA with potential for PHB production. When looking at the fate of propionyl-CoA, the cumulative ΔG going to 3-hydroxyvaleryl-CoA formation (with the final reaction to 3HV assumed) is −35.72 KJ/mol, whereas going to 3-hydroxybutanoyl-CoA (with the final reaction to PHB assumed) through the MCC pathway and phosphoenolpyruvate is −109.9058 KJ/mol. This suggests that even if propionyl-CoA is present, there is more thermodynamic drive toward PHB production than 3HV production.

The findings from MDF analysis support the experimental results mentioned earlier. For strain sPS015, the increase in PHB suggests that there is in fact some internal sourcing of propionyl-CoA happening, however, ultimately it is more thermodynamically favorable to go toward PHB rather than 3HV. This is also seen in strain sPS014 where formation of propionyl-CoA both in the presence and absence of propionate leads to an increase in PHB yield. The production of 3HV by sPS014, but not sPS015 may be explained by substrate availability. As was already discussed in the context of PHB, β-ketothiolase, the first enzyme in both the PHB and 3HV pathways, is thermodynamically unfavorable in the forward direction. This means that a high amount of propionyl-CoA would be needed to push the reaction in the forward direction toward 3-ketovaleryl-CoA (Alsiyabi et al., 2021). This is a condition that appears to be met when propionate is fed into the cell and improved upon with the overexpression of *pct*, but does not appear to be satisfied by the amount of propionyl-CoA that can be internally sourced.

Combined, these results show that the interconnectedness of propionyl-CoA and acetyl-CoA in *P. sacchari*’s metabolic network create competition between PHB and 3HV production, with PHB production being more favorable. Previous work attempting to block propionyl-CoA from being recycled back to acetyl-CoA using targeted gene knockouts was not entirely successful (Pereira et al., 2009), yet those methods combined with either strategy shared here may yield better 3HV results. We suggest that further exploration into additional pathways that may be competing for propionyl-CoA would be a worthwhile endeavor toward development of *P. sacchari* as a bioproduction platform for high 3HV fraction PHBV.

## 4. Conclusion

The findings in this study demonstrate the use of promoters pLAC and BBaJ23 104 for control of metabolic genes for increased PHB and 3HV yield. Maximum yield increases were seen in strain sPS012, which increased PHB by 162%, and sPS014 in the absence of propionate, which increased 3HV by 145%. While these improvements show promise, perhaps it was the failed strategies that were more informative for future work. Our hypothesis that strategies to increase conversion of fed-in propionate and to allow for internal sourcing of propionyl-CoA would improve 3HV production was not supported. Yet, these results combined with postulation from the MDF analysis allow for a better understanding of factors controlling the fate of PHB and 3HV precursors. Future work to fully elucidate and separate propionyl-CoA from PHB synthesis could drastically improve 3HV yields in *P. sacchari* and allow for more tunable control of PHBV production.

## Acknowledgments

We would like to thank Drs. Bara Altartouri and You Zhou for their assistance with the TEM work done at the Microscopy Research Core Facility of the Center for Biotechnology, which is partially funded by the Nebraska Center for Integrated Biomolecular Communication COBRE grant (NIH P20 GM113126) and the Nebraska Research Initiative. All figures were made in Biorender (https://BioRender.com).

