## Supplemental Materials for "Heterologous expression of critical pathway genes leads to complex pattern of increased yield of bioplastic precursors in *Paraburkholderia sacchari* ITP 101"

Supplementary Materials

Supplementary Table 1. Sequence of genetic parts used in this study including genes, terminators, and promoters.

| **Genetic Parts** | **Sequence** | **source** |
|---|---|---|
| gfpUV | atgagtaaaggagaagaacttttcactggagttgtcccaattcttgttgaattagatggtgatgttaatgggcacaaattttctgtcagtggagagggtgaaggtgatgcaacatacggaaaacttacccttaaatttatttgcactactggaaaactacctgttccgtggccaacacttgtcactactttctcttatggtgttcaatgcttttcccgttatccggatcacatgaaacggcatgactttttcaagagtgccatgcccgaaggttatgtacaggaacgcactatatctttcaaagatgacggggaactacaagacgcgtgctgaagtcaagtttgaaggtgatacccttgttaatcgtatcgagttaaaaggtattgattttaaagaagatggaaacattctcggacacaaactggagtacaactataactcacacaatgtatacatcacggcagacaaacaaaagaatggaatcaaagctaacttcaaaattcgccacaacattgaagatggctccgttcaactagcagaccattatcaacaaaatactccaattggcgatggccctgtccttttaccagacaaccattacctgtccacacaatctgcccttcgaaagatcccaacgaaaagcgtgaccacatggtccttcttgagtttgtaactgctgctgggattacacatggcatggatgagctctacaaataa | Plasmid: pBbB7a-GFP (Lee et al., 2011) |
| phaA | atgactgacgttgtcatcgtatccgccgcccgcaccgcggtcggcaaatttggcggctcgctggccaagatcccggcgccggaactgggtgccgtggtcatcaaggccgcgctggaacgcgccggcgtcaagccggagcaggtgagcgaagtcatcatgggccaggtgctgaccgccggttcgggccagaaccccgcacgccaggccgcgatcaaggccggcttgccggcgatggtgccggccatgaccatcaacaaggtgtgcggctcgggcctgaaggccgtgatgctggccgccaacgcgatcatggcgggcgacgccgagatcgtggtggccggcggccaggaaaacatgagcgccgccccgcacgtgctgccgggctcgcgcgacggcttccgcatgggcgatgccaagctggtcgacaccatgatcgtcgacggcctgtgggacgtgtacaaccagtaccacatgggcatcaccgccgagaatgtggccaaggaatacggcatcacgcgcgaggcgcaggatgagctcgcggtcggctcgcagaacaaggccgaagccgcgcagaaggccggcaagtttgacgaagagatcgtcccggtgctgatcccgcagcgcaagggcgacccggtggccttcaagaccgacgagttcgtgcgccagggcgccacgctggacagcatgtccggcctcaagcctgccttcgacaaggccggcacggtgaccgcggccaacgcctcgggcctgaacgacggcgccgccgcggtggtggtgatgtcggcggccaaggccaaggaactgggcctgaccccgctggccacgatcaagagctacgccaacgccggtgtcgatcccaaggtgatgggcatgggcccggtgccggcctccaagcgcgccctgtcgcgcgccgagtggaccccgcaagacctggacctgatggagatcaacgaggcctttgccgcgcaggcgctggcggtgcaccagcagatgggctgggacacctccaaggtcaatgtgaacggcggcgccatcgccatcggccatccgatcggcgcgtcgggctgccgtatcctggtgacgctgctgcacgagatgaagcgccgtgacgcgaagaagggcctggcctcgctgtgcatcggcggcggcatgggcgtggcgctggcagtcgagcgcaaataa | *C. necator* gDNA |
| bktb | atgacgcgtgaagtggtagtggtgagcggtgtccgtaccgcgatcgggacctttggcggcagcctgaaggatgtggcgccggcggagctgggcgcgctggtggtgcgcgaggcgctggcgcgcgcgcaggtatcgggcgacgatgtcggccacgtggtgttcggcaacgtgatccagaccgagccgcgcgacatgtatctgggccgcgtcgcggccgtcaacggcggggtgacggtcaatgcccccgcgctgaccgtgaaccgcctgtgcggttcgggcctgcaggccatcgtcagcgccgcgcagactatcctgctgggcgataccgacgtcgccatcggcggtggcgcggaaagcatgagccgcgcgccgtacctggcgcccgcggcccgctggggcgcacgcatgggcgatgccggcctggtcgacatgatgctgggcgcgctgcacgatcccttccatcgcatccacatgggcgtgaccgccgagaacgtcgccaaggaatacgacatctcgcgcacgcagcaggacgaggccgcgctggaatcgcaccgccgcgcttcggccgcgatcaaggccggttacttcaaggaccagatcgtcccggtggtgagcaagggccgcaagggcgatgtcaccttcgacaccgacgagcacgtgcgtcatgacgccaccatcgacgacatgaccaagctcaggccggtcttcgtcaaggaaaatggcacggtcacggccggcaattcctcgggcctgaacgacgccgccgccgcggtggtgatgatggagcgcgccgaagccgagcgccgcggcctgaagccgctggcccgtctggtgtcgtacggccatgccggcgtggacccccaagaccatgggcatcggcccggtgccggcgaccaagatcgcgctggagcgcgccggcctgcaggtgtcggacctggacgtaatcgaagccaacgaagcctttgccgcgcaggcgtgcgcggtgaccaaggcgctcggcctggatccggccaaggtcaacccgaacggctcgggcatctcgctgggccacccgatcggcgccaccggtgccctgatcacggtgaaggcgctgcatgagctgaaccgcgtgcagggccgctacgcgctggtgacgatgtgcatcggcggcgggcagggcattgccgccatcttcgagcgtatctga | *C. necator* gDNA |

| | | |
|---|---|---|
| phaB | atgACTCAGCGCATTGCGTATGTGACCGGCGGCATGGGTGGTATCGG<br>AACCGCCATCTGCCAGCGGCTGGCCAAGGATGGCTTTCGTGTGGTG<br>GCCGGCTGCGGCCCCAACTCGCCGCGCCGTGAAAGGTGGCTCGAGC<br>AGCAGAAGGCACTCGGCTTTGACTTCGTTGCTTCTGAAGGCAACGT<br>GGCTGACTGGGACTCGACCAAGACCGCATTCGACAAGGTCAAGGCC<br>GAGGTCGGCGAGGTCGATGTGCTGATCAACAACGCCGGCATCACCC<br>GCGACGTGGTGTTCCGCAAGATGACCCGTGCCGACTGGGACGCGGT<br>GATCGACACTAACCTGACCTCGCTGTTCAACGTCACCAAGCAGGTG<br>ATCGACGGCATGGCCGACCGCGGGTGGGGCCGTATCGTCAACATCT<br>CGTCGGTGAACGGGCAGAAGGGCCAGTTCGGCCAGACCAACTACTC<br>CACCGCCAAGGCCGGCCTACACGGCTTCACCATGGCGCTGGCGCAG<br>GAAGTGGCGACCAAGGGTGTGACGGTCAACACCGTTTCGCCGGGCT<br>ATATCGCCACCGACATGGTCAAGGCGATCCGCCAGGACGTGCTCGA<br>CAAGATCGTCGGGACGATCCCGGTCAAGCGCCTGGGCGAGCCGGA<br>AGAGATCGCTTCGATCTGCGCCTGGCTGTCGTCGGACGAGTCCGGT<br>TTCTCGACCGGCGCTGACTTCTCGCTCAACGGCGGCCTGCATATGG<br>GCtga | *C. necator* gDNA |
| phaC | atggcgaccggcaaaggcgcggcagcttccactcaggaaggcaagtcccaaccattcaagttcacgccgggg<br>ccattcgatccagccacatggctggaatggtcccgccagtggcagggcactgaaggcaacggccacgcggcc<br>gcgtccggcattccgggcctggatgcgctggcaggcgtcaagatcgagccggcgcagctgggtgatatccag<br>cagcgttacatgaaggacttctcagccctgtggcaggccatggccgagggcaaggccgaggccaccgggcc<br>gctgcacgaccggcgcttcgccggcgacgcgtggcgcaccaacctgccataccgcttcgctgccgcgttctac<br>ctgctcaatgcgcgcgccttgaccgagctggccgatgctgttgaggccgatgccaagacgcgccagcgcatcc<br>gctttgcgatctcgcaatgggtcgatgcgatgtcgcccgccaacttcctcgccacgaatcccgaggcgcagcgc<br>ctgctgatcgagtcgggcggcgaatcgctgcgtgccggcgtgcgcaacatgatggaagacctgacgcgcggc<br>aagatctcgcagaccgacgagagcgcgtttgaggtcggccgcaatgtcgcggtgagcgaaggcgccgtagtc<br>ttcgagaacgaatacttccagctgttgcagtacaagccgctgaccgacaaggtgcatgcgcgcccgctgctgat<br>ggtgccgccgtgcatcaacaagtactacatcctggacctgcagccggagagctcgctggtgcgtcatgtggtgg<br>agcaggggcatacggtgttcctggtgtcgtggcgcaatccggacgccagcatggctggcagcacctgggacg<br>actacatcgagcacgcggccatccgcgccatcgaagtcgcgcgcgacatcagcggccaggacaagatcaac<br>gtgctcggcttctgcgtgggcggcaccattgtgtcgactgcgctggcggtgatggccgcgcgcggccagcacc<br>cggctgccagcgtcacgctgctgaccacgctgctggactttgccgacaccggcatcctcgacgtctttgtcgac<br>gagggccatgtgcagctgcgcgaggccacgctgggcggcgccgccggcgcgccgtgcgcgctgctgcgcg<br>gccttgagctggccaataccttctcgttcctgcgcccgaacgacctggtgtggaactacgtggtcgacaactacc<br>tgaagggcaacacgccggtgccgttcgacctgctgttctggaacggcgacgccaccaacctgccggggccttg<br>gtactgctggtacctgcgccacacctacctgcaggacgagctcaaggtgccgggcaagctgactgtgtgcggc<br>gtgcccgtggacctggccagcatcgacgtgccgacctacatctacggctcgcgcgaagaccatatcgtgccat<br>ggaccgcggcctatgcctcgaccgcgctgctggcgaacaagctgcgcttcgtgctgggtgcgtcgggccatat<br>cgccggtgtgatcaacccgccggccaagaacaagcgcagccactggaccaacgatgcgctgccggagtcgc<br>cgcagcaatggctggccggcgccatcgagcatcacggcagctggtggccggactggaccgcatggctggcc<br>gggcaggccggcgcgaaacgcgccgcgcccgccaactatggcaatgcgcgctatcgcgcaatcgaacccg<br>cgcctgggcgatacgtcaaagccaaggcatga | *C. necator* gDNA |

| | | |
|---|---|---|
| sbm operon | GATTAATGACCAACGAAATTAGGTTTACGTTTTTCGAGGAAAGCGT<br>TCATCCCTTCCTGGTAATCTTCGCTGTCATACACCGCGCGGCGCATC<br>CCCTGAATACGTTCAAATTCATCGGAGTTCATGGTGTGTGCTTCGCC<br>CAGTACACGCAGCTCTTCTTTGATAACGGCAATGGCTAACGGCGCT<br>TTCTCAGAGATGTGGTGCGCCATTTGTAAGGTGAAATCTTCCAGTTC<br>TTCCACTTCCACAACATGGTTGAGGATGCCGACAGCCAGCGCGCGC<br>TGGGCGGTGATTGGCGAAGCGGTAAAAATCAGCTCTTTGACAATGT<br>GGAAGCCCGCGTCGCGGGTCAGGTTGTGAATGCCGACCAGGTTATA<br>CGGGACGCCGAGGTTTACAGGCGTCATTGAGAAGGTTGAGGTACTG<br>GCGGCGATGATCAGATCGGAACTCATGATCATTTCAAATGCGCCAC<br>CCCAAACACTACCTTCCACCATCGAAATGATCGGTTTCGGGAATTTT<br>TGGATCATGCGGGTGATTTGACGCAATGGATCATCATAGGAGAGCG<br>GATCGCGACCGCCAGACGGCAGTTCGTGAATATCGTGACCTGCGGA<br>GAAGACTTTGGATCCACTCGGTGCGCGCAAAATGATACAGCGAATT<br>TCCGGCCGGTTGAGATCGCTTAACGCCTGCATAAGATCATCAATAA<br>AGACTTTACTTAAGGCATTAAGTTTTCGGCCATAGTTAAACTCAATG<br>ACCGCCACTTTGTTGATAGTGACAACGTTAACATACTGATAAGACA<br>TAAAAATTCCTTTAATCAAAATATTGCGTCTGGATAAATTCACTGAG<br>CTGCCGCAGGCCGGTGCGCGGTGAGAGCGTATTGTTTTTGACCGCT<br>AAAAGCGTCTGGCGGTAATAGCGATCGAAATCTTCATTCGCGAACA<br>GGTGATTCAGTACTTCTTCTTCGGTCTGCTTACGCAGCCATTCCACC<br>GATTGTTGTTGCCGCACTTGTTGTAAACGACCACTGGCAGTTAGCGC<br>GGTTTTGAAGTCGATGATGGCGTGCCAGATCTCATCGATTCCACGTT<br>TTTCCAGTGCGCTACAAGTCAGAACCCGTGGCTGCCATTCGTCGTAT<br>TTACGTCGCAGAATATGCAGGGCACTCTCGTACATATGCCGGGCAA<br>TGGCGACATTGGTATGGTTATCGCCATCGTCTTTGTTGATAACGATC<br>AGATCAGCCACTTCCATCAGCCCTTTTTTAATGCCCTGCAGATCATC<br>GCCGCCACCGGCAATTTGCAACGAGATAAAACAGTCCACCATGCGG<br>GCGACTTCTGTTTCCGACTGCCCGACGCCAACCGTTTCGACAATCAC<br>TACGTCATAACCCGCTGCTTCGCATAACAGCATTAATTCCCGCGCTC<br>GCTGACTGGCACCGCCCAGATGACCGGAGGATGGTACCGGGCGAA<br>TAAACGCCGCTTCGGCACGCGCCAGGTCATTCATGCGGGTTTTATCC<br>CCGAGAATGCTACCGCCAGTGACCGGGCTGCTGGGATCGACCGCAA<br>TAACCGCGACCTTTAATCCCTCTCGAATCAACAACATGCCAAAGGC<br>CTCAAGAAAGGTACTTTTCCCCGCGCCGGGGGTGCCGGTAACGCCC<br>AGTCGCAGGGTGTTACCGCAGTACGGCATAATGGCATCAAGCAGCT<br>GCGTACTTAGTGCCTGATGACGCGGGTGACGGCTTTCCACCAGCGT<br>CATGGCCTGGGCGAGTGTGGCACGCTCACCCTGACGTAAGCGGCGA<br>ATACTTTCTGCCAGCGTGGCTTCATTAATCATGATGCTGGCTTATCA<br>GATTCAGTACGTCGCGCACACTGTCGAGCATAGGTGTACCTGGACC<br>ATAAATCGCCGCCACGCCGCGCTCTTGCAGGAAGGCGTAATCCTGC<br>GGCGGAATGACGCCACCCGCGACCACGCAGATATCTTCGCGTCCCC<br>ATTTTTTCAGCGCTTCGACCAGTTCCGGGATCAGCGTTTTATGACCG<br>GCAGCCAGTGAGGATGCGCCCACTACGTGAACGTCGTTTTCTACGG<br>CCAGGCGGGCGATCTCTTCAGGTGTAGAGAACATCGGGCTTAAATC<br>TACGTCGAAACCGAGATCGGAATAGGCGCTGGCGATCACTTTCGCG<br>CCGCGATCGTGTCCATCCTGGCCCATCTTAGCGATCAGAATGCGCG<br>GGCGACGACCATTGTCGGCAAGGAACTGCTCCGTTTGCGCAACAAT<br>GGCATCGAACTCGGAGGCCGATTTCTCAGACTGATGATAGCTTTGC<br>GCAATCACGCCGGTAACACACTGGCTTGGCACCAGATAACGGTCGA<br>AAGCGACTTCCAGCGCATCGGAAATTTCACCCAGGGTGGCGCGAAC<br>GCGAGCGGCATTAACAGCGGCAGCCAGCAGGTTTTCGTTATGCTGT<br>GCGGCGTGAGTCAGGGCGTTCAACGCGGCGGTTACGGCGGCATCAT<br>CACGGGTGGCGCGAATGCGTTCCAGCGAAGCAATTTGCTCGTTACG<br>CACCATCACGTTGTCGATCTCAAGTACATCGGTTTCGTCTTCGTGAT<br>CCAGTTTGTACTTGTTGACACCAACGATGACACGCTTGCCCTGGTCG<br>ATCAGCGACTGTTCGCGCGCTGAGGCCTCTTCGATCATTCGTTTTGG<br>CAGACCTGCTTCGATCGCTTTCGCCATGCCACCGGCTTCGTCGATCT<br>GTTGGATAATAGCTCTGGCTTGTTTGACGATTTGATCGGTCAGCGAC<br>TCAATGTAATAGGATCCGGCCAGTGGATCGACGGTGCGGCAGAGTT<br>CTGATTCTTCCTGGATGATGATCTGGGTGTTGCGGGCAATGCGTGCT<br>GAGAAATCGGTAGGCAAACCAAGCGCTTCGTCAAAGACGTTGGTAT | *E. coli* gDNA |

| | |
|---|---|
| | GCAGTGACTGAGTACCGCCCAGCGTCGCAGCCAGCGCTTCAATGGT<br>GGTGCGGATAACGTTGTTATACGGATCCTGTTCAGTCAGGCTCCAG<br>CCTGAGGTCTGGCAGTGGGTACGCAGCGCCAGTGATTTCGGGTCCT<br>GTGCGCCAAATCCACTGACCGCTTCGCTCCATAAATAACGTGCCGC<br>ACGCAACATGGCGACGTTCATAAACAGATCCATGCCGATGCCGAAG<br>AAGAACGACAGGCGAGGAGCGAAGTCATCAATTTTCAGTCCGGCA<br>GAGATTGCTGCTTTGATGTACTCAATCCCATCAGCGAGCGTAAATG<br>CTACCTGCTGCACGCAGTTGGCACCCGCTTCACCCATGTGGTAACC<br>GCTGATACTGATGGTATTAAATCGCGGCATGTTGCCGGAACACCAG<br>GCGATGATGTCGGCGATAATGCGCATTGACGGTTTTGGTGGGTAAA<br>TATAGGTGTTGCGGCAGAGGTACTCTTTGAGAATATCGTTTTGAATG<br>GTGCCGGTCAGTTTATCAGGTGTAACACCTTGCTCTTCTGCGGCGAC<br>GATATAAAACGCCAGTACTGGTAGCACTGCGCCATTCATGGTCATC<br>GAAACCGACATTTTATCCAGCGGGATCTGGTCGAACAGGACTTTCA<br>TATCTTCCACGGTGTCGATAGCGACGCCCGCTTTGCCGACGTCGCCC<br>GCCACGCGCGGGTTATCGGAGTCGTAGCCACGGTGGGTGGCAAGGT<br>CAAACGCAACGGAAAGACCTTTTTGCCCGGCGGCCAGGTTACGGCG<br>ATAAAAAGCGTTGGACTCTTTTGCTGTTGAAAAACCAGCATACTGA<br>CGGATGGTCCACGGTTGGGCGGTATACATAGTGGCACGCGGGCCAC<br>GAACGTAGGGCGGCAAACCAGGAAGGGTACCTGTCACCTCCAGATT<br>ATCGAGATCGGCTTCGGTATACAGCGGCTTGATGGCGATCCCTTCC<br>GCGGTTTGATGAACCAGCGAGTCGACAGTTTTCTCCCGACGGCTCA<br>ATTCCTTGTTGGCAAGCTGTTGCCACTCCTGCACGTTAGACAT |

| | | |
|---|---|---|
| pct | ATGAAGGTGATCACCGCACGCGAAGCGGCGGCACTGGTGCAGGAC<br>GGCTGGACCGTGGCCAGCGCGGGCTTTGTCGGCGCCGGCCATGCCG<br>AGGCCGTGACCGAGGCGCTGGAGCAGCGCTTCCTGCAGAGCGGGCT<br>GCCGCGCGACCTGACGCTGGTGTACTCGGCCGGGCAGGGCGACCGC<br>GGCGCGCGCGGCGTGAACCACTTCGGCAATGCCGGCATGACCGCCA<br>GCATCGTCGGCGGCCACTGGCGCTCGGCCACGCGGCTGGCCACGCT<br>GGCCATGGCCGAGCAGTGCGAGGGCTACAACCTGCCGCAGGGCGT<br>GCTGACGCACCTATACCGCGCCATCGCCGGCGGCAAGCCCGGCGTG<br>ATGACCAAGATCGGCCTGCACACCTTCGTCGACCCGCGCACCGCGC<br>AGGATGCGCGCTACCACGGCGGCGCCGTCAACGAGCGCGCGCGCC<br>AGGCCATTGCCGAGGGCAAGGCATGCTGGGTCGATGCGGTCGACTT<br>CCGCGGCGACGAATACCTGTTCTACCCGAGCTTCCCGATCCACTGC<br>GCGCTGATCCGCTGCACCGCGGCCGACGCCCGCGGCAACCTCAGCA<br>CCCATCGCGAAGCCTTCCACCATGAGCTGCTGGCGATGGCGCAGGC<br>GGCCCACAACTCGGGCGGCATCGTGATCGCGCAGGTGGAAAGCCTG<br>GTCGACCACCACGAGATCCTGCAGGCCATCCACGTGCCCGGCATCC<br>TGGTCGACTACGTGGTGGTCTGCGACAACCCCGCCAACCACCAGAT<br>GACGTTTGCCGAGTCCTACAACCCGGCCTACGTGACGCCATGGCAA<br>GGCGAGGCAGCGGTGGCCGAAGCGGAAGCGGCGCCGGTGGCTGCC<br>GGCCCGCTCGACGCGCGCACCATCGTGCAGCGCCGTGCGGTGATGG<br>AACTGGCGCGCCGTGCGCCGCGCGTGGTCAACCTGGGCGTGGGCAT<br>GCCGGCAGCGGTCGGCATGCTGGCGCACCAGGCCGGGCTGGACGG<br>CTTCACGCTGACCGTCGAGGCCGGCCCCATCGGCGGCACGCCCGCG<br>GATGGCCTCAGCTTCGGTGCCTCGGCCTACCCGGAGGCGGTGGTGG<br>ATCAGCCCGCGCAGTTCGATTTCTACGAGGGCGGCGGCATCGACCT<br>GGCCATCCTCGGCCTGGCCGAGCTGGATGGCCACGGCAACGTCAAT<br>GTCAGCAAGTTCGGCGAAGGCGAGGGCGCATCGATTGCCGGCGTCG<br>GCGGCTTTATCAACATCACGCAGAGCGCGCGCGCGGTGGTGTTCAT<br>GGGCACGCTGACGGCGGGCGGGCTGGAAGTCCGCGCCGGCGACGG<br>CGGCCTGCAGATCGTGCGCGAAGGCCGCGTGAAGAAGATCGTGCCT<br>GAGGTGTCGCACCTGAGCTTCAACGGGCCCTATGTGGCGTCGCTCG<br>GCATCCCGGTGCTGTACATCACCGAGCGCGCGGTGTTCGAGATGCG<br>CGCTGGCGCAGACGGCGAAGCCCGCCTCACGCTGGTCGAGATCGCC<br>CCCGGCGTGGACCTGCAGCGCGACGTGCTCGACCAGTGCTCGACGC<br>CCATCGCCGTTGCGCAGGACCTGCGCGAAATGGATGCGCGGCTGTT<br>CCAGGCCGGGCCCCTGCACCTGTAA | *C. necator* gDNA |
| rrnBT1 terminator | caaataaaacgaaaggctcagtcgaaagactgggcctttcgttttatctgttgtttgtcggtgaacgctctc | (Mutalik et al., 2013) |
| pLAC | tttacactttatgcttccggctcgtatgttg | pBBR1MCS-2 (Kovach et al., 1995) |
| BBa_J23 100 | ttgacggctagctcagtcctaggtacagtgctagc | Anderson promoters (Anderson, 2008) |
| BBa_J23 104 | ttgacagctagctcagtcctaggtattgtgctagc | Anderson promoters |
| BBa_J23 107 | tttacggctagctcagccctaggtattatgctagc | Anderson promoters |
| BBa_J23 115 | tttatagctagctcagcccttggtacaatgctagc | Anderson promoters |

Supplementary Table 2. Primer pairs used in this study.

| Primer pairs | Target | Template DNA | | Sequence |
|---|---|---|---|---|
| pbackbone | Kanamycin resistance, PBBR1 ori, pLAC promoter | pBBR1MCS-2 | forward | ttgttatccgctcacaattccaca |
| | | | reverse | cagtttgctcaggctctccccgt |
| pbackbone2 | Kan resistance, PBBR1 ori, pLAC promoter, rrnBT1 terminator | PS-gfp-plac | forward | ttgttatccgctcacaattccaca |
| | | | reverse | atccaaactcgagtaaggatct |
| pgfp-plac | *gfpUV*, rrnBT1terminator | AS-Plac(c)-GFPuv-amp | forward | gagagcctgagcaaactggcctcag gcatttgagaagca |
| | | | reverse | attgtgagcggataacaatttcatttca gaattcaaaag |
| pPS100 | Kan resistance, PBBR1 ori, *gFPuv*, rrnBT1 terminator | PS-gfp-plac | forward | ctaggtacagtgctagcGCCTG GGGTGCCTAATGAGTG |
| | | | reverse | gactgagctagccgtcaaTGTGG AATTGTGAGCGGATAA C |
| pPS104 | Kan resistance, PBBR1 ori, *gFPuv*, rrnBT1 terminator | PS-gfp-plac and PS-pha(A/B/C)/bktb-plac | forward | ctaggtattgtgctagcGCCTGG GGTGCCTAATGAGTG |
| | | | reverse | gactgagctagctgtcaaTGTGG AATTGTGAGCGGATAA C |
| pPS107 | Kan resistance, PBBR1 ori, *gFPuv*, rrnBT1 terminator | PS-gfp-plac | forward | ctaggtattatgctagcGCCTGG GGTGCCTAATGAGTG |
| | | | reverse | ggctgagctagccgtaaaTGTGG AATTGTGAGCGGATAA C |
| pPS115 | Kan resistance, PBBR1 ori, *gFPuv*, rrnBT1 terminator | PS-gfp-plac | forward | cttggtacaatgctagcGCCTGG GGTGCCTAATGAGTG |
| | | | reverse | ggctgagctagctataaaTGTGG AATTGTGAGCGGATAA C |
| pphaAplac | *phaA* | *C. necator* gDNA | forward | attgtgagcggataacaacaggttcc ctcccgtttccattgaaaggactac |
| | | | reverse | ccttactcgagtttggatcactccttga ttggctcttcgttatcgtcg |
| pphaBplac | *phaB* | *C. necator* gDNA | forward | attgtgagcggataacaaCGAAG AGCCAATCAAGGAGTG |
| | | | reverse | ccttactcgagtttggatCTGGCT GCACCGCAATACG |
| pphaCplac | *phaC* | *C. necator* gDNA | forward | attgtgagcggataacaaggttcgaa tagtgacggcagagagacaatc |
| | | | reverse | ccttactcgagtttggatgtagtcctttc aatggaaacgggagggaac |
| pbktbplac | *bktb* | *C. necator* gDNA | forward | attgtgagcggataacaactgaaacg ggttgattaggtaaaagtacgctc |
| | | | reverse | ccttactcgagtttggatcagcgtgca gaggttcttcgtcagc |

| | | | | |
|---|---|---|---|---|
| psbm | *sbm operon* | *E. coli* gDNA | forward | ccttactcgagtttggatATGTCTAACGTGCAGGAGTGGC |
| | | | reverse | attgtgagcggataacaaGATTAATGACCAACGAAATTAGGTTTAC |
| ppct | *pct* | *C. necator* gDNA | forward | ccttactcgagtttggatAATGACAGGATTACAGGTGCAG |
| | | | reverse | attgtgagcggataacaaATGAAGGTGATCACCGCACG |

Supplementary Table 3. Reaction KEGG IDs, equations, KEGG equations, and estimated $\Delta G$ for *P. sacchari* model used for MDF analysis.

| KEGG rxn ID | Equation | KEGG equation | Estimated $\Delta G$ |
|---|---|---|---|
| R00299 | ATP + D-Glucose <=> ADP + D-Glucose 6-Phosphate | C00002 + C00031 <=> C00008 + C00092 | -20.4 |
| R00771 | D-Glucose 6-phosphate <=> D-Fructose 6-phosphate | C00092 <=> C00085 | 2.6 |
| R00756 | ATP + D-Fructose 6-phosphate <=> ADP + D-fructose 1,6-bisphosphate | C00002 + C00085 <=> C00008 + C00354 | -17.8 |
| R01068 | D-Fructose 1,6-bisphosphate <=> Glycerone phosphate + D-Glyceraldehyde 3-phosphate | C00354 <=> C00111 + C00118 | 23.2 |
| R01061 | D-Glyceraldehyde 3-phosphate + Orthophosphate + NAD+ <=> 3-Phospho-D-glyceroyl phosphate + NADH + H+ | C00118 + C00009 + C00003 <=> C00236 + C00004 + C00080 | 1.2 |
| R01512 | ADP + 3-Phospho-D-glyceroyl phosphate <=> ATP + 3-Phospho-D-glycerate | C00008 + C00236 <=> C00002 + C00197 | -19.5 |
| R01518 | 3-Phospho-D-glycerate <=> 2-Phospho-D-glycerate | C00197 <=> C00631 | 4.5 |
| R00658 | 2-Phospho-D-glycerate <=> Phosphoenolpyruvate + H2O | C00631 <=> C00074 + C00001 | -3.8 |
| R00200 | ADP + Phosphoenolpyruvate <=> ATP + Pyruvate | C00008 + C00074 <=> C00002 + C00022 | -25 |
| R00209 | Pyruvate + CoA + NAD+ <=> Acetyl-CoA + CO2 + NADH + H+ | C00022 + C00010 + C00003 <=> C00024 + C00011 + C00004 + C00080 | -34.2 |
| R00351 | Acetyl-CoA + H2O + Oxaloacetate <=> Citrate + CoA | C00024 + C00001 + C00036 <=> C00158 + C00010 | -38.8 |
| R01325 | Citrate <=> cis-Aconitate + H2O | C00158 <=> C00417 + C00001 | 8.5 |
| R01324 | Citrate <=> Isocitrate | C00158 <=> C00311 | 6.8 |
| R01900 | Isocitrate <=> cis-Aconitate + H2O | C00311 <=> C00417 + C00001 | 1.6 |
| R01899 | Isocitrate + NADP+ <=> Oxalosuccinate + NADPH + H+ | C00311 + C00006 <=> C05379 + C00005 + C00080 | 15.7 |
| R00267 | Isocitrate + NADP+ <=> 2-Oxoglutarate + CO2 + NADPH + H+ | C00311 + C00006 <=> C00026 + C00011 + C00005 + C00080 | 5.4 |
| R00268 | Oxalosuccinate <=> 2-Oxoglutarate + CO2 | C05379 <=> C00026 + C00011 | -10.3 |
| R08549 | 2-Oxoglutarate + CoA + NAD+ <=> Succinyl-CoA + CO2 + NADH + H+ | C00026 + C00010 + C00003 <=> C00091 + C00011 + C00004 + C00080 | -28.2 |

| | | | |
|---|---|---|---|
| R00405 | ADP + Orthophosphate + Succinyl-CoA <=> ATP + Succinate + CoA | C00008 + C00009 + C00091 <=> C00002 + C00042 + C00010 | 1.2 |
| R02164 | Quinone + Succinate <=> Hydroquinone + Fumarate | C15602 + C00042 <=> C15603 + C00122 | 64.85 |
| R01082 | Fumarate + H2O <=> (S)-Malate | C00122 + C00001 <=> C00149 | -3.4 |
| R00342 | (S)-Malate + NAD+ <=> Oxaloacetate + NADH + H+ | C00149 + C00003 <=> C00036 + C00004 + C00080 | 26.5 |
| R00431 | GTP + Oxaloacetate <=> GDP + Phosphoenolpyruvate + CO2 | C00044 + C00036 <=> C00035 + C00074 + C00011 | 13.3 |
| R00925 | ATP + Propanoate + CoA <=> AMP + Diphosphate + Propanoyl-CoA | C00002 + C00163 + C00010 <=> C00020 + C00013 + C00100 | -1.1 |
| R00931 | Propanoyl-CoA + Oxaloacetate + H2O <=> 2-Methylcitrate + CoA | C00100 + C00036 + C00001 <=> C02225 + C00010 | -40.4 |
| R04424 | 2-Methylcitrate <=> (Z)-But-2-ene-1,2,3-tricarboxylate + H2O | C02225 <=> C04225 + C00001 | -2.9 |
| R04425 | (Z)-But-2-ene-1,2,3-tricarboxylate + H2O <=> (2S,3R)-3-Hydroxybutane-1,2,3-tricarboxylate | C04225 + C00001 <=> C04593 | 2.9 |
| R00409 | (2S,3R)-3-Hydroxybutane-1,2,3-tricarboxylate <=> Pyruvate + Succinate | C04593 <=> C00022 + C00042 | 5.9 |
| R00238 | 2 Acetyl-CoA <=> CoA + Acetoacetyl-CoA | 2C00024 <=> C00010 + C00332 | 25 |
| R01977 | Acetoacetyl-CoA + NADPH + H+ <=> (R)-3-Hydroxybutanoyl-CoA + NADP+ | C00332 + C00005 + C00080 <=> C03561 + C00006 | -14.3 |
| RPHV1 | acetyl coA + propionyl coA <=> CoA + 3-ketovaleryl-coA | C00024+CPHV1 <=> C00010 + CPHV2 | 29.53 |
| RPHV2 | 3-ketovaleryl coA + NADPH + H+ <=> HV CoA + NADP+ | CPHV2 + C00005 + C00080 <=> CPHV3 + C00006 | 0.6972 |
| R00833 | Succinyl-CoA <=> (R)-Methylmalonyl-CoA | C00091 <=> C01213 | 8.4 |
| R02765 | (R)-Methylmalonyl-CoA <=> (S)-Methylmalonyl-CoA | C01213 <=> C00683 | -0.2 |
| R00923 | (S)-Methylmalonyl-CoA <=> Propanoyl-CoA + CO2 | C00683 <=> C00100 + C00011 | -16.6 |
